# Selective Treg recruitment to bone remodeling niches is required for digit tip regeneration

**DOI:** 10.64898/2026.05.04.722813

**Authors:** Chunyan Fu, Char Wynter, Emily A. Polk, Kailin R. Mesa

## Abstract

Adult mammals have limited capacity for tissue regeneration, where most injuries resolve through fibrotic scarring rather than functional tissue restoration^1–4^. Studies in regenerative vertebrate species, including amphibians, teleost fish, reptiles and mammals, have established that the innate immune system plays instructive roles in regeneration^5–11^, yet the role of adaptive immune cells and how the immune response distinguishes regenerative from non-regenerative injuries, remain poorly understood. The mouse digit tip provides a rare mammalian model of complete multi-tissue regeneration where distal amputation through the terminal phalanx (P3) triggers complete multi-tissue regrowth, whereas a more proximal amputation of the same bone results in fibrotic scarring^12–16^. Using an intravital multiphoton imaging approach capable of longitudinally tracking bone remodeling and immune cells in live mice^17^, we find that regulatory T cells (Tregs) are selectively recruited to regenerating but not scarring digit tips. Tregs localize first to sites of osteoclast-mediated bone resorption and persist at the bone surface when an expanding stromal progenitor pool, known as the blastema, initiates digit regrowth. Acute depletion of Tregs impairs bone resorption and subsequent bone regrowth. Mice lacking T and B cells or CD4+ and CD8+ T cells show similar bone remodeling defects, suggesting a dominant role for Tregs within the adaptive immune compartment in promoting mammalian digit tip regeneration. Treg depletion impairs regeneration through an IL-10-independent mechanism, pointing to a non-canonical effector program. Lastly, pharmacological blockade of the chemokine receptor CXCR4 reduces Treg recruitment to the bone compartment, diminishes bone-associated macrophage accumulation, and attenuates bone degradation in regenerative amputations. Together, these findings identify Tregs as essential regulators of bone remodeling during mammalian digit tip regeneration.

## Main Text

Adult mammals repair most injuries through fibrotic scarring rather than functional tissue regeneration, a response that limits recovery across diverse organs from cardiac muscle, lung and spinal cord^1–4^. In contrast, several non-mammalian vertebrates accomplish true epimorphic regeneration, defined as the complete structural and functional restoration of amputated tissues^6^. Classic examples of epimorphic regeneration include the axolotl limb and zebrafish fin^5,18^. Epimorphic regeneration requires two conserved hallmarks: blastema formation, a transient structure of stromal progenitor cells, and an intact immune system, as perturbation of innate immune cells, including macrophages, severely impairs this process^7–11^. These observations have established that the immune system not only responds to general tissue damage but also has an essential role in functional regeneration of lost tissues.

Despite this progress, a fundamental question remains unresolved: how does the immune system differentially respond to regenerative and scarring injuries? This question is difficult to address in classical non-mammalian models because these organisms often regenerate without a natural scarring comparison after injury. Moreover, recent work has demonstrated that lymphocyte-deficient axolotls and zebrafish regenerate amputated appendages with normal kinetics, patterning, and skeletal outcomes^19,20^, suggesting that adaptive immunity may be dispensable for epimorphic regeneration in highly regenerative vertebrates. By contrast, lymphocytes, in particular regulatory T cells (Tregs), have been shown to improve injury repair in multiple mammalian tissues, including skeletal muscle, skin, lung, and thymus^21–24^, suggesting the mammalian system has co-opted lymphoid components, including Tregs, to support regeneration in ways not observed in species with more robust intrinsic regenerative capacity. Addressing both questions requires a mammalian model that naturally produces both regenerative and scarring (non-regenerative) responses to injury in similar tissue contexts.

The terminal or third phalanx (P3) of the mouse digit provides precisely such a model. Distal amputation produces a reproducible regenerative response, while proximal amputation of the same bone results in scarring^12,13,15^. This dichotomy within a single bone element provides two injuries with similar systemic immune environments, yet divergent regenerative outcomes. Prior work has established that osteoclast-mediated bone degradation is an obligate and rate-limiting first step in the regenerative program: bone resorption creates a pocket at the resulting bone surface that enables subsequent formation of the blastema progenitor pool. Bone degradation initiates at approximately 5 days post-amputation (DPA 5) and is complete by DPA 10, at which point the blastema begins to form and expand until DPA 13; differentiation and patterning then proceed progressively with substantial bone regrowth apparent at both DPA 21 and 28^14,16^. The non-regenerative (proximal) amputation fails to initiate this program, but when precisely the two injury responses diverge, and what immune signals underlie these distinct tissue remodeling outcomes, remain unclear.

Understanding the cellular dynamics of digit tip regeneration has been constrained by several technical challenges, including highly dynamic and temporally compressed transition points, architecturally complex tissues, and limited cell numbers that make it difficult to adequately resolve cellular events by conventional histological sectioning or flow cytometry. Furthermore, while macrophage requirement has been established in digit tip regeneration^8^, whether and how adaptive immune cells participate remains largely unexplored. To overcome these limitations, we developed a longitudinal intravital two-photon imaging platform that tracks bone structure, immune cells, and stromal progenitors simultaneously in live mice across the full regenerative program. Using this platform in combination with fluorescent reporter mouse lines, genetic and antibody-mediated cellular depletion strategies, and pharmacological perturbations, we identify regulatory T cells as selectively recruited and functionally essential participants in mammalian digit tip regeneration.

## Intravital imaging reveals early divergent bone remodeling

To establish a longitudinal imaging system capable of resolving bone architecture and immune cell behavior in the intact digit, we adapted an intravital multiphoton microscopy approach that we have used previously to study skin macrophage dynamics^17^. Label-free second harmonic generation (SHG) from fibrillar collagen within bone provided a longitudinal structural readout of bone organization without exogenous labeling^25,26^. To validate that SHG faithfully reports bone mineral content across the regenerative window, we co-labeled bone with two independent mineralizing dyes, Calcein and Alizarin red S, and confirmed close co-localization with SHG signal at multiple time points (Extended Data Fig. 1a, b). We then performed serial imaging of both distal (regenerative, P3 R) and proximal (scarring, P3 NR) amputations at DPA 0, 7, 14, 21, and 28, paired with micro-CT analysis at the same time points (Figure 1a).

**Figure 1.**
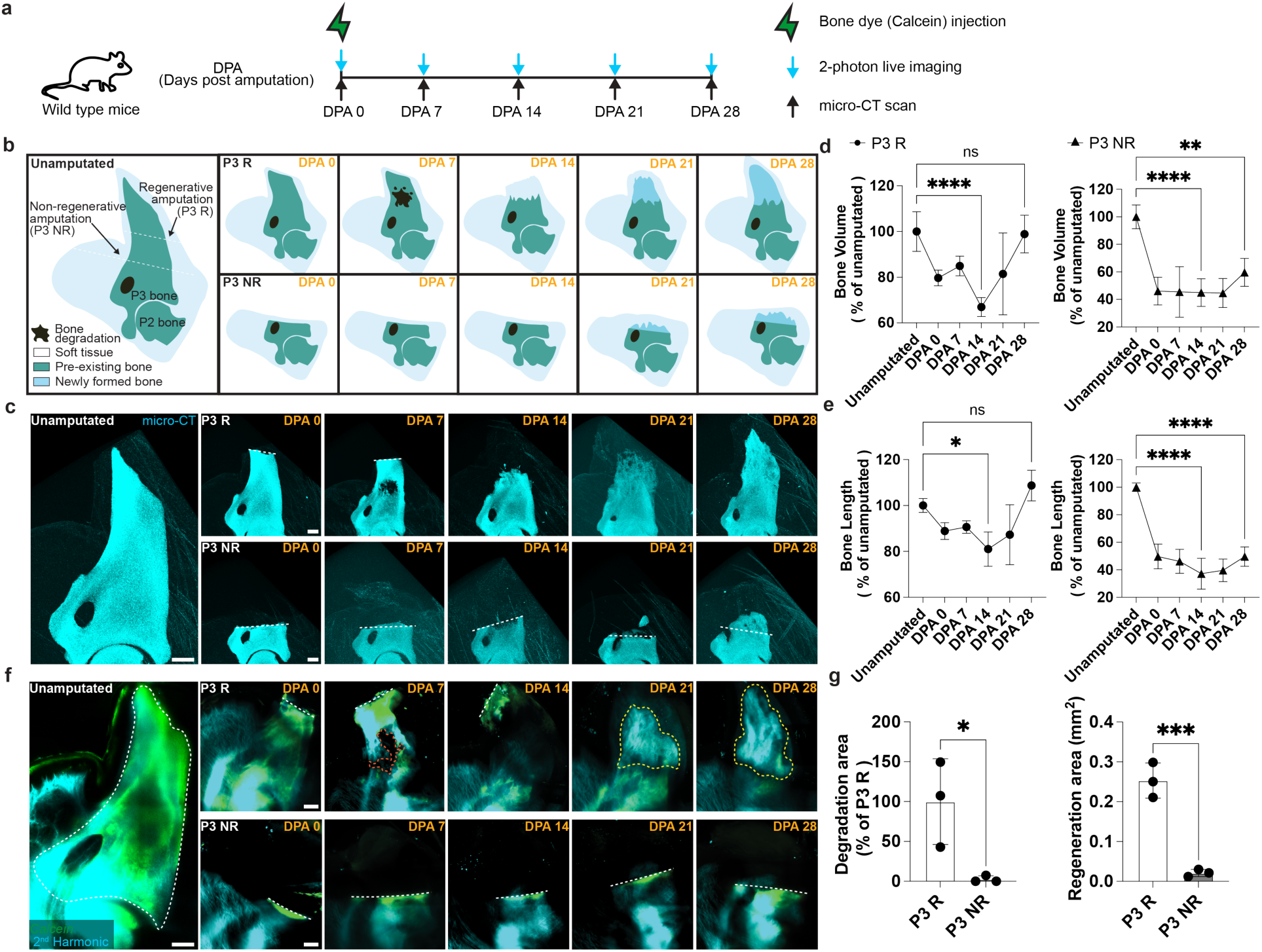
Intravital imaging reveals early divergent bone remodeling in regenerative and scarring digit amputations. a, Timeline showing Calcein bone dye injection at DPA 0 and serial two-photon and micro-CT imaging at unamputated, DPA 0, 7, 14, 21, and 28. **b,** Schematic cross-sections of unamputated, P3 R, and P3 NR digit tips across the time course. Dark fill, bone resorption cavity; dark blue, pre-existing bone; light blue, newly formed bone; gray, soft tissue. **c,** Representative micro-CT images of unamputated, P3 R (top), and P3 NR (bottom) digit tips across the time course. Dashed white outlines indicate the amputation plane. **d,** Quantification of bone volume for P3 R (left, n = 3-6 mice) and P3 NR (right, n = 3-6 mice) across the time course (mean ± SD; *p < 0.05; ns, not significant; two-way ANOVA with Tukey’s test). **e,** Quantification of bone length for P3 R (left, n = 3-6 mice) and P3 NR (right, n = 3-6 mice) (mean ± SD; *p < 0.05; ns; two-way ANOVA with Tukey’s test). **f,** Representative two-photon images showing SHG bone signal (cyan) and Calcein (green) in unamputated, P3 R (top rows) and P3 NR (bottom rows) digit tips at DPA 0, 7, 14, 21, and 28. Dashed white lines (unamputated), P3 bone; dashed orange outlines (DPA 7), degradation area; dashed yellow outlines (DPA 21, 28), regeneration area; dashed straight white outlines indicate the amputation plane. **g,** Quantification of degradation area (left) and regeneration area (right) in P3 R (n = 3 mice) versus P3 NR (n = 3 mice) (mean ± SD; *p < 0.05, ***p < 0.001; unpaired Student’s t-test). Scale bar, 150 μm.

In P3 R digits, a reproducible and stereotyped sequence of bone remodeling was apparent. SHG imaging revealed progressive erosion of the distal bone surface and formation of a degradation cavity by DPA 7, and subsequent in-filling with new mineralized matrix from DPA 14 through DPA 28 (Figure 1b, c, f, Supplementary Video 1). Micro-CT quantification of P3 R and P3 NR digits found a significant decrease at DPA 14 in both bone volume and length when compared to unamputated controls. By DPA 28, P3 R digits showed no significant difference to unamputated controls, while P3 NR digits remained significantly decreased in bone volume and length (Figure 1d, e). Utilizing our two-photon intravital imaging approach, we assessed bone degradation and regeneration at DPA 7 and 21, respectively, in live mice. Quantification of bone degradation and regeneration areas confirmed that these processes were both significantly restricted to P3 R digits (Figure 1g). Selective bone degradation in P3 R digits at DPA 7 was also apparent by micro-CT imaging (Supplementary Video 2). Together, these data validate our intravital imaging platform and reveal that non-regenerative digit amputations fail to initiate the earliest step in the regenerative program, bone degradation, identifying this as a critical early divergence point between the two injury responses.

## Tregs are selectively recruited to regenerating digit tips

The divergence in bone remodeling between P3 R and P3 NR digit tips raised the question of whether immune cell composition differs between conditions at the time of this earliest divergence. Tregs have been shown to promote tissue repair across multiple mammalian contexts^21–24^, and single-cell transcriptomic analysis of regenerating digit tips has identified CD3+ T cells expressing Receptor Activator of Nuclear Factor kappa B Ligand (RANKL) at DPA 8, a known driver of osteoclast activation and bone degradation^27^. These observations motivated us to directly assess whether Tregs are selectively recruited to the bone remodeling niche during regeneration. To do so, we performed longitudinal intravital imaging in Forkhead box P3 (Foxp3) reporter mice (*Foxp3-RFP*), enabling direct visualization of Foxp3+ Tregs across the full regenerative time course (Figure 2a).

**Figure 2.**
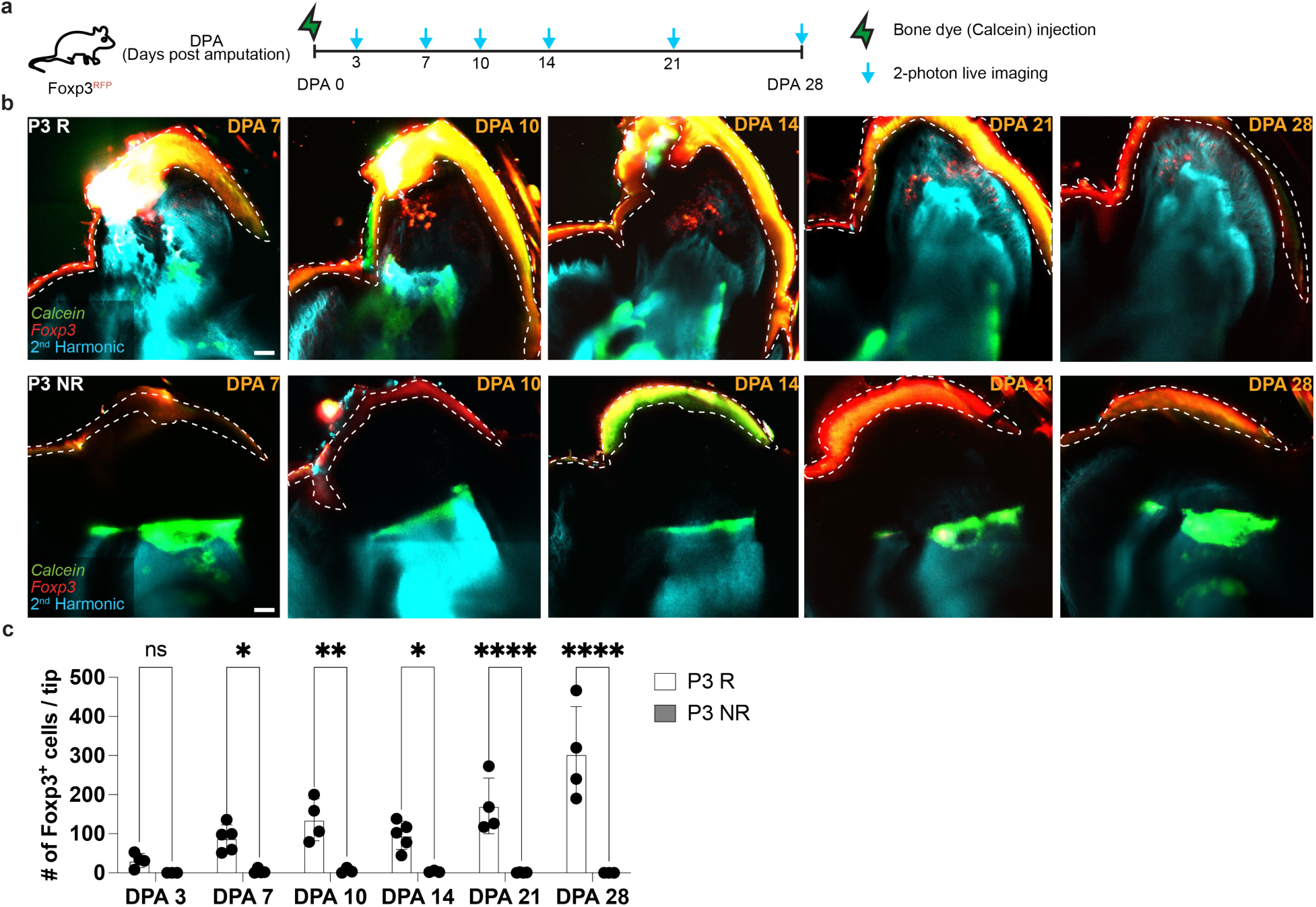
Tregs are selectively recruited to regenerating digit tips. a, Schematic timeline for live imaging of *Foxp3-RFP* reporter mice: Calcein injection at DPA 0 and serial two-photon imaging at DPA 3, 7, 10, 14, 21, and 28. **b,** Representative two-photon images showing Foxp3-RFP+ Tregs (red) with SHG (cyan) and Calcein (green) in P3 R (top) and P3 NR (bottom) digit tips at DPA 7, 10, 14, 21, and 28. Dashed white line: digit tip boundary; color within: autofluorescence. **c,** Quantification of Foxp3+ Treg number per mouse in P3 R (white bars, n = 4-5 mice per time point) and P3 NR (gray bars, n = 3-4 mice per time point ) across the time course (mean ± SD; ns, *p < 0.05, **p < 0.01, ****p < 0.0001; two-way ANOVA with Tukey’s test). Scale bar, 100 μm.

Serial imaging from DPA 3 through DPA 28 revealed a selective accumulation of Foxp3+ Tregs in P3 R but not P3 NR digit tips (Figure 2b, c). Tregs were rare or absent in both conditions at DPA 3 but began accumulating in P3 R digits by DPA 7 as bone degradation commenced. Treg numbers in P3 R digits increased progressively through DPA 28, while P3 NR digits showed no significant Treg enrichment at any time point when compared to DPA 3. To assess whether this selectivity reflected a shift in T cell composition rather than overall T cell recruitment, we performed flow cytometric profiling of P3 R and P3 NR digit tips at DPA 7 (Extended Data Fig. 2). Overall CD3+ T cell proportions were not significantly different between conditions, indicating that the selective Treg accumulation we observed represents a change in T cell composition rather than a global increase in T cell infiltration. Together, these data demonstrate that Treg recruitment is concomitant with the regenerative injury program and is not a general response to digit tip amputation.

## Tregs localize to sites of bone resorption and blastema formation

To determine where Tregs localize relative to the cellular events of digit tip regeneration, we performed dual reporter intravital imaging in *Foxp3-RFP*; Colony-stimulating factor 1 receptor (Csf1r) reporter mice (*Csf1r-EGFP*), enabling simultaneous visualization of Tregs (red), macrophages (including osteoclasts) (green), and bone SHG (cyan) (Figure 3a). At DPA 5, Tregs were already detectable within and immediately adjacent to the P3 bone surface, prior to overt sites of bone degradation (Figure 3b, c). By DPA 7, when the bone degradation cavity was well established, the majority of Tregs were found within the degradation zone itself, near bone-associated macrophages at the degrading bone surface (Figure 3d, e, Supplementary Video 3).

**Figure 3.**
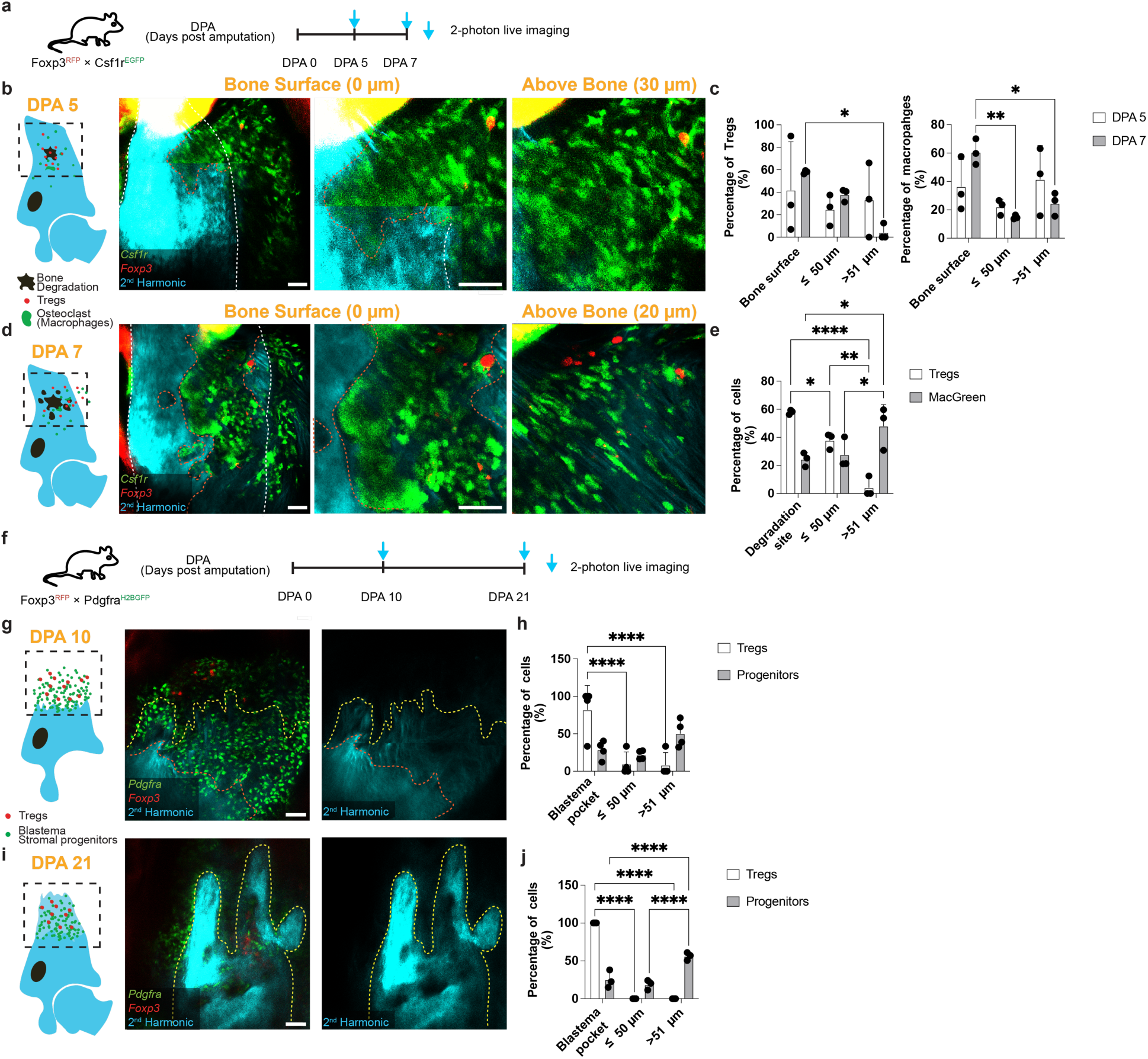
Tregs concentrate at sites of bone resorption and blastema formation. a, Timeline of imaging protocol in *Foxp3-RFP*; *Csf1r-EGFP* dual-reporter mice (DPA 5, and 7). **b, d** Schematics (left) and representative two-photon images (right) at DPA 5 (**b**) and DPA 7 (**b**) showing Tregs (red), macrophages (green), and bone (cyan) with panels showing at degrading bone surface (center) and above bone degradation area (right). Dashed white lines outline P3 bone, dashed orange lines outline degradation areas. **c,** Quantification of the percentage of Tregs and macrophages located at the bone surface, within 50 μm, or beyond 51 μm at DPA 5 (n = 3 mice) and DPA 7 (n = 3 mice) (mean ± SD; *p < 0.05, **p < 0.01; two-way ANOVA with Tukey’s test). **e,** Quantification of the percentage of Tregs and macrophages located in the degradation site, within 50 μm, or beyond 51 μm at DPA 7 (n = 3 mice) (mean ± SD; *p < 0.05, **p < 0.01, ****p < 0.0001; two-way ANOVA with Tukey’s test). **f,** Schematic timeline of imaging protocol in *Foxp3-RFP*; *Pdgfrα-H2BGFP* dual-reporter mice (DPA 10, and 21). **g, i,** Schematics (left) and representative two-photon images (right) at DPA 10 (**g**) and DPA 21 (**i**) showing Tregs (red) and PDGFRα+ blastema stromal progenitors (green). Dashed orange lines outline degradation areas; dashed yellow lines outline regeneration areas. **h, j,** Quantification of the percentage of Tregs and blastema progenitor cells inside the bone cavity, within 50 μm, or beyond 51 μm at DPA 10 (n = 4 mice) and DPA 21 (n = 3 mice) (mean ± SD; ****p < 0.0001; two-way ANOVA with Tukey’s test. Scale bar, 50 μm.

To assess Treg spatial dynamics during the later regrowth phase of regeneration, we performed imaging in *Foxp3-RFP*; Platelet-Derived Growth Factor Receptor Alpha (*Pdgfrα*) reporter mice (*Pdgfrα-H2BGFP)*, co-labeling Tregs (red), blastema stromal progenitor cells (green), and bone SHG (cyan) (Figure 3f). From DPA 10 through DPA 21, as the PDGFRα+ progenitor cell pool expanded distally at the base of the degraded bone surface, Tregs were found distributed within the blastema pocket throughout this period (Figure 3g–j, Supplementary Video 4). This temporally ordered recruitment pattern suggests that Tregs participate in both the degradation and regrowth phases of digit tip regeneration.

## Tregs are required for digit tip regeneration

To test whether Tregs are functionally required for digit tip regeneration, we used *Foxp3-DTR* mice, in which diphtheria toxin (DT) administration specifically depletes Foxp3+ cells^28^. DT was administered beginning at DPA −1 and repeated at regular intervals throughout the imaging period (Figure 4a). Flow cytometric analysis of digit tips confirmed efficient and sustained Treg depletion in DT-treated *Foxp3-DTR* mice compared to control *Foxp3-DTR* mice (Extended Data Fig. 3a, b). Body weight was not significantly different between groups across the experiment, suggesting that the depletion protocol did not cause overt systemic inflammation (Extended Data Fig. 3c). Longitudinal SHG imaging revealed a significant reduction in bone degradation area at DPA 7 and impaired bone regeneration at DPA 21 in Treg-depleted mice compared to DT-treated WT mice (Figure 4b–e). Micro-CT analysis at DPA 28 confirmed significantly reduced P3 bone volume and length in DT-treated *Foxp3-DTR* mice compared to controls (Figure 4f, g). These results demonstrate a requirement for Tregs in both the early osteoclast-driven bone resorption and the subsequent blastema-driven bone regrowth phases of digit tip regeneration.

**Figure 4.**
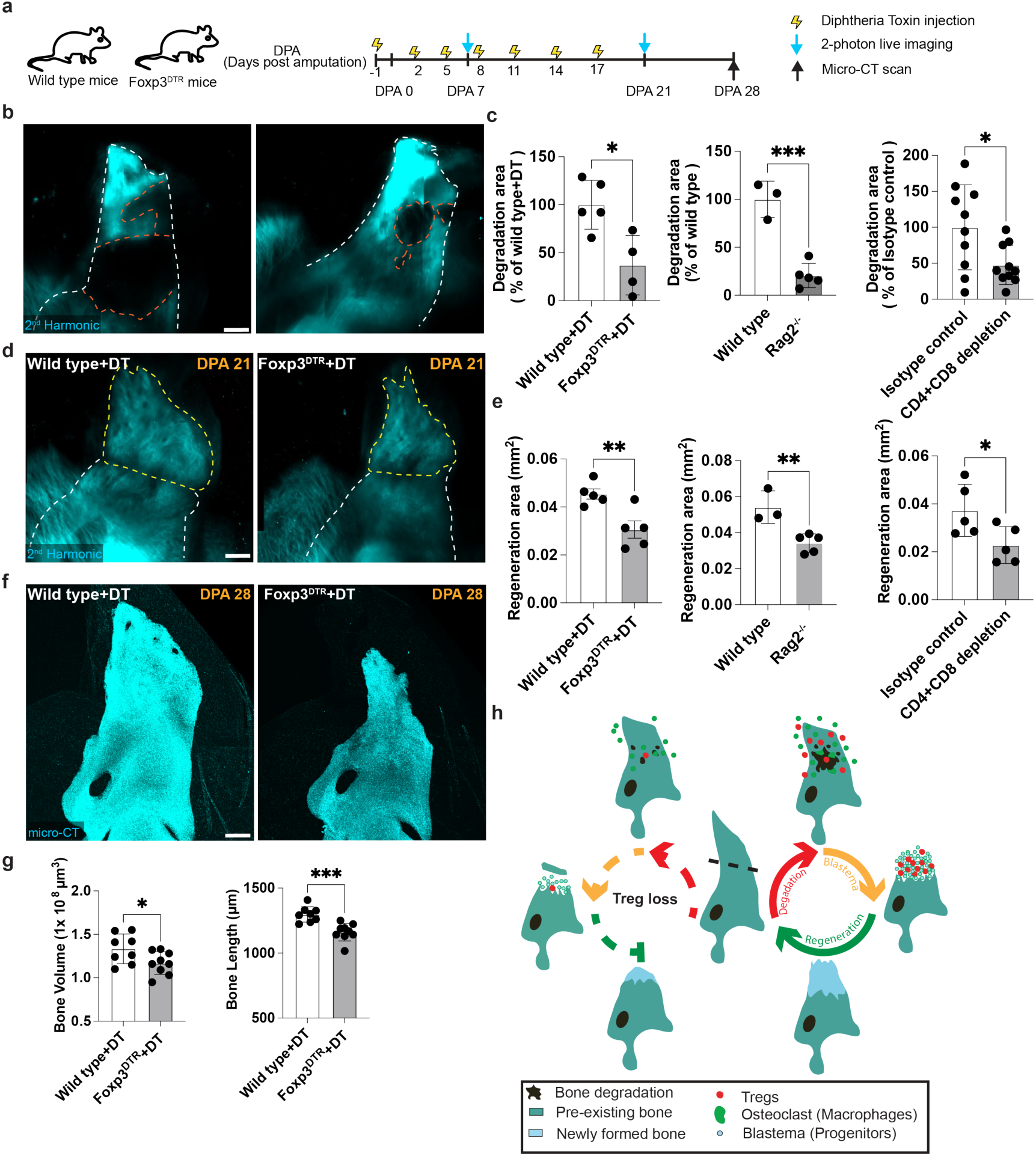
Tregs are required for digit tip bone regeneration. a, Schematic timeline of diphtheria toxin (DT) administration (beginning at DPA −1, repeated at DPA 2, 5, 8, 11, 14, and 17) and imaging sessions in Wild-type and *Foxp3-DTR* mice. **b,** Representative SHG images at DPA 7 from Wild-type + DT (left) and *Foxp3-DTR* + DT (right) mice. Dashed white lines outline P3 bone; dashed orange lines outline degradation area. **c,** Quantification of bone degradation area at DPA 7 across three independent perturbation strategies: Wild-type + DT (n = 5 mice) vs (n = 5 mice) *Foxp3-DTR* + DT (left), Wild-type (n = 3 mice) vs *Rag2*^-/-^ (n = 5 mice) (center), and isotype control (n = 5 mice) vs anti-CD4 + anti-CD8 antibody depletion (n = 5 mice) (right) (mean ± SD; *p < 0.05, ***p < 0.001; unpaired Student’s t-test). **d,** Representative SHG images at DPA 21 from Wild-type + DT (left) and *Foxp3-DTR* + DT (right). Dashed white lines outline P3 bone; dashed yellow lines outline regeneration area. **e,** Quantification of bone regeneration area at DPA 21 across the same three perturbation strategies: Wild-type + DT (n = 5 mice) vs (n = 4 mice) *Foxp3-DTR* + DT (left), Wild-type (n = 3 mice) vs *Rag2*^-/-^ (n = 5 mice) (center), and isotype control (n = 10 mice) vs anti-CD4 + anti-CD8 antibody depletion (n = 10 mice) (right) (mean ± SD; *p < 0.05, **p < 0.01; unpaired Student’s t-test). **f,** Representative micro-CT images at DPA 28 from Wild-type + DT (left) and *Foxp3-DTR* + DT (right). **g,** Quantification of P3 bone volume (left) and bone length (right) at DPA 28 in Wild-type + DT mice (n = 8 mice) vs *Foxp3-DTR* + DT (n =9 mice) (mean ± SD; *p < 0.05, ***p < 0.001; unpaired Student’s t-test). Scale bar, 150 μm. **h,** Working model illustrating the consequence of Treg loss: impaired bone degradation leads to failure of blastema formation and regeneration.

Given that prolonged Treg depletion can unleash systemic autoimmune inflammation through uncontrolled effector T cell responses^28,29^, we complemented the *Foxp3-DTR* data with two independent T cell perturbation strategies. First, we performed distal P3 amputations in Recombination Activating Gene 2 (Rag2)-deficient (*Rag2*^-/-^) mice, which lack mature T and B cells. Intravital imaging at DPA 7 and DPA 21 revealed reduced bone degradation and regeneration in *Rag2*^-/-^ mice compared to Wild-type controls (Figure 4c, e, Extended Data Fig. 4c, d). Second, we performed acute antibody-mediated depletion of CD4+ and CD8+ T cells in Wild-type mice. Efficient depletion was confirmed by flow cytometry at DPA 0 and DPA 7 (Extended Data Fig. 5a-c). Imaging of depleted mice revealed a similar pattern of impaired bone degradation (DPA 7) and regeneration (DPA 21) (Figure 4c, e, Extended Data Fig. 5d, e). Quantification across all three perturbation strategies confirmed a consistent and significant reduction in both bone degradation at DPA 7 and bone regeneration at DPA 21 (Figure 4c, e). Notably, recent studies have reported that adoptive transfer of NK and CD4+ T cells can exert cytotoxic effects on digit tip progenitor cells, and that Tregs may protect progenitor cells from this cytotoxicity^30,31^. Our data extend this model by demonstrating that Tregs also support the earlier bone resorption phase required to establish the permissive environment for blastema formation, indicating that Tregs are required at multiple stages in the regenerative program.

To explore whether canonical Treg-mediated immunosuppression through Interleukin-10 (IL-10) underlies this pro-regenerative function^32^, we performed distal amputations in IL-10^-/-^ mice. In contrast to the impairment observed after Treg depletion, IL-10^-/-^ mice showed no significant reduction in bone degradation area at DPA 7, bone regeneration area at DPA 21, or final bone volume and length at DPA 28 (Extended Data Fig. 6). These data indicate that Tregs support digit tip regeneration through mechanisms other than canonical IL-10-mediated immunosuppression^33^.

## CXCR4 mediates Treg homing to the bone niche

Tissue recruitment of Tregs is known to rely on multiple chemokine signaling pathways^34^; however, recent work has highlighted an essential role for C-X-C motif chemokine receptor 4 (CXCR4) / C-X-C motif chemokine ligand 12 (CXCL12) signaling in recruiting Tregs to sites of tissue regeneration^35,36^. Therefore, we next evaluated the role of CXCR4 signaling in Treg recruitment during digit tip regeneration. We administered a CXCR4 antagonist (AMD3100) daily beginning at DPA 0 and assessed consequences for Treg recruitment and bone remodeling at DPA 7 (Figure 5a). Bone degradation was significantly reduced in CXCR4 antagonist-treated mice compared to PBS controls (Figure 5b, c), consistent with a role for CXCR4-dependent Treg homing in enabling productive bone resorption. This reduction in bone degradation was accompanied by a decrease in the total number of Foxp3+ Tregs detected in the digit at DPA 7 (Figure 5d, e). At the same point, there was a marked decrease in co-localization between macrophages and sites of degradation, without a significant change in total number (Figure 5f–h), suggesting impaired or reduced osteoclast activity.

**Figure 5.**
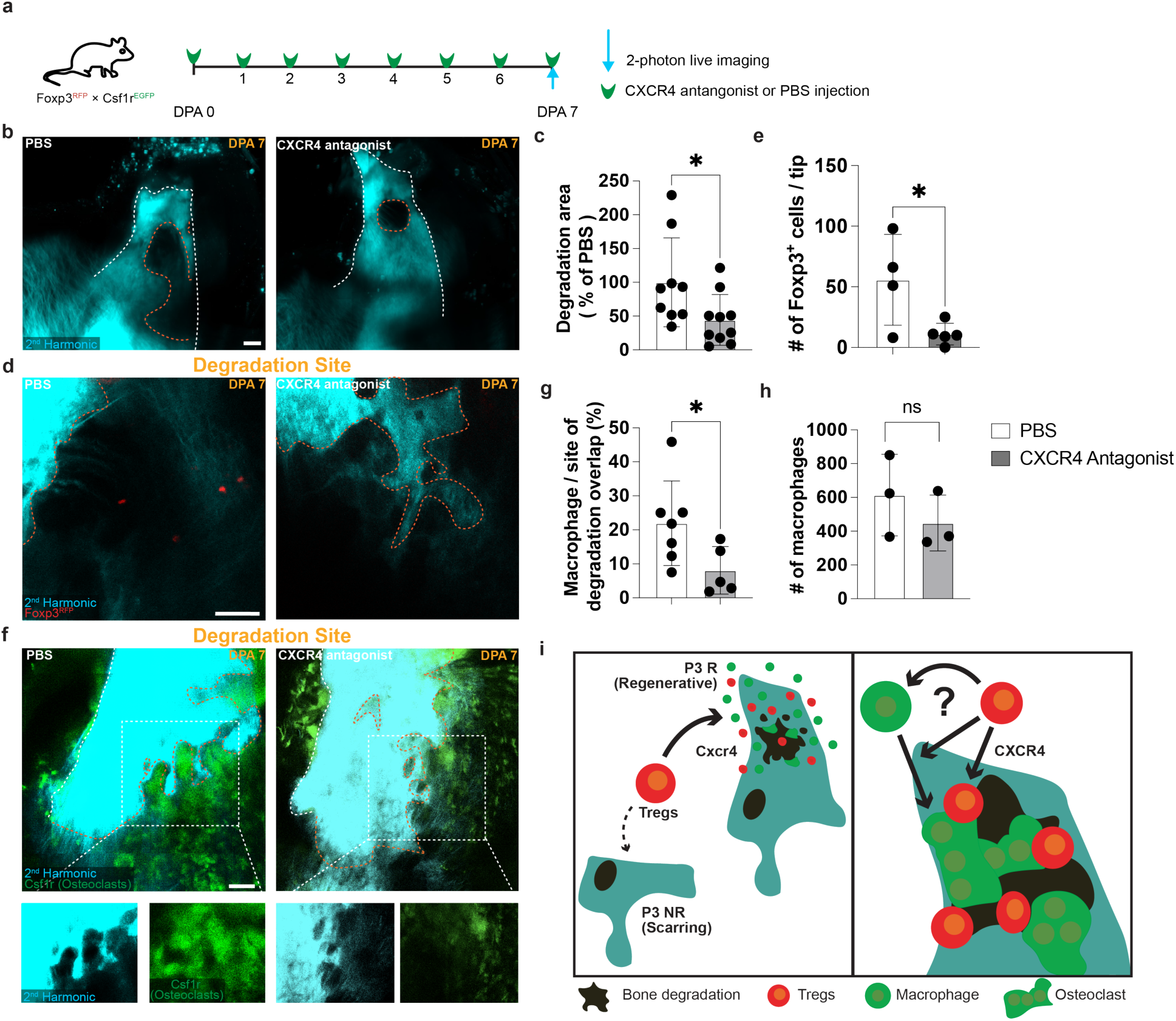
CXCR4 signaling mediates Treg recruitment to the bone degradation niche. a, Schematic timeline of CXCR4 antagonist (AMD3100) or PBS administration (daily, DPA 0 through DPA 7) and two-photon imaging at DPA 7 in *Foxp3-RFP*; *Csf1r-EGFP* dual-reporter mice. **b,** Representative SHG images showing bone degradation cavity in PBS (left) and CXCR4-antagonist (right) conditions. Dashed white lines outline P3 bone; dashed orange lines outline degradation area. **c,** Quantification of bone degradation area at DPA 7 (n = 8 mice for PBS and 9 mice for AMD3100) (mean ± SD; *p < 0.05; unpaired Student’s t-test). **d,** Representative two-photon images of *Foxp3-RFP*+ Tregs (red) in PBS-treated (left) and CXCR4-antagonist-treated (right) digit tips at DPA 7. Dashed orange lines outline degradation area. **e,** Quantification of total Foxp3+ Treg number per digit tip (left, *p < 0.05) in PBS (n =3 mice) versus CXCR4 antagonist-treated mice (n =5 mice) (mean ± SD; unpaired Student’s t-test). **f,** Representative two-photon images of macrophages (green) and bone (cyan) in PBS and CXCR4 antagonist conditions at DPA 7. Dashed orange lines outline degradation area. **g, h,** Quantification of macrophage/degradation site overlap percentage (**g**) and macrophage number (**h**) in PBS (n =3-7 mice) versus CXCR4 antagonist-treated mice (n =3-5 mice) (mean ± SD; unpaired Student’s t-test). **i,** Model: CXCR4-mediated Treg recruitment promotes macrophage localization to the bone resorption niche in regenerative (P3 R) amputations. Absence of Tregs in the non-regenerative (P3 NR) condition correlates with macrophage exclusion from the bone surface and failure to initiate bone degradation. Scale bar, 50 μm.

These data support a model in which CXCR4 signaling is required, for efficient Treg recruitment to the bone degradation niche in regenerating digit tips. Impaired Treg homing under CXCR4 blockade correlates with reduced macrophage localization to the degradation sites, consistent with a role for Tregs in supporting osteoclast activation at the resorption front (Figure 5i). Our data are consistent with recent work identifying CD3+ T cells as a source of RANKL, a well-known activator of osteoclasts, in regenerating digit tips at DPA 8 by single-cell transcriptomics^22^. Treg-specific conditional deletion of CXCR4 will be required to formally establish whether this homing requirement is cell-intrinsic. Key open questions include whether CXCR4 acts directly on Tregs, the identity of the CXCL12-producing cell(s) in the regenerating digit, and whether additional chemokine receptors cooperate with CXCR4 in Treg homing.

## Discussion

The immune system plays instructive roles in tissue regeneration across vertebrates. Macrophages are required for blastema formation and appendage regrowth in non-mammalian models^5,18^, and their depletion similarly impairs mouse digit tip regeneration^8^, establishing an evolutionarily conserved innate immune requirement for epimorphic regeneration. Yet two fundamental questions have remained unresolved: whether the adaptive immune system participates in this process as well, and how immune responses are differentially coordinated to support regeneration rather than fibrotic scarring after injury. The mouse digit tip is uniquely suited to address both, as distal and proximal amputations of the same bone produce complete regeneration and fibrotic scarring, respectively, within a shared systemic immune environment^12^. Using longitudinal intravital two-photon microscopy, we find that regulatory T cells are selectively recruited to regenerating but not scarring digit tips, localize sequentially to sites of active bone resorption and blastema formation, and are required for successful tissue regrowth. These findings identify Tregs as the first adaptive immune population shown to be selectively recruited to and required for regenerative bone remodeling and establish selective Treg recruitment as a key axis by which the immune system distinguishes regenerative from scarring injury.

The requirement for Tregs manifests at two temporally and spatially distinct stages of the regenerative program. Tregs localize first to the osteoclast-active bone resorption front at DPA 5 through DPA 7 and then persist within the expanding blastema as stromal progenitor cells fill the degradation cavity from DPA 10 onward. Acute Treg depletion impairs bone degradation at DPA 7 as well as bone regrowth at DPA 21, establishing that Tregs are required at both stages. The same bone remodeling defect was observed in *Rag2^−/−^* mice and in Wild-type mice depleted of CD4+ and CD8+ T cells, establishing a dominant role for Tregs rather than other lymphocyte populations within the adaptive immune compartment. Importantly, IL-10-deficient mice regenerate with normal kinetics and bone volumes, ruling out canonical Treg-mediated immunosuppression as the operative mechanism. Pharmacological blockade of CXCR4, a chemokine receptor required for Treg homing to several tissue niches^34–36^, reduces Treg recruitment to the digit, decreases macrophage localization within the bone degradation cavity, and impairs bone resorption. This ordered relationship positions CXCR4-dependent Treg recruitment upstream of productive macrophage localization at the resorption front, identifying Treg homing as a rate-limiting step in the innate-adaptive coordination required for bone remodeling.

Collectively, these findings establish that mammalian digit tip regeneration requires coordinated innate and adaptive immune activity, with Tregs and macrophages acting in sequence to enable bone resorption and blastema formation. Tregs act through a non-canonical, IL-10-independent mechanism and are recruited via CXCR4 to the resorption niche, where their presence correlates with productive macrophage positioning at the degrading bone surface. The co-requirement for both innate and adaptive immune populations in mammals, but not in highly regenerative non-mammalian vertebrates^19,20^, suggests that the mammalian immune system has co-opted adaptive immune components to enable regenerative tissue remodeling that would otherwise resolve as fibrotic repair.

## Acknowledgements

We are grateful to all members of the Mesa laboratory for valuable discussions and critical reading of the manuscript. We thank the Flow Cytometry Core at Princeton University, and specifically C.J. Decoste and J. Garcia, for technical support with FACS. We are grateful to D.V. Potapenko for assistance with the Zeiss Xradia 630 Versa 3D X-ray microscope at the Imaging and Analysis Center, Princeton University. We thank J.F. Brooks II for providing IL-10^-/-^ mice. This work was supported by internal startup funds from Princeton University.

## Author Contributions

C.F. and K.R.M. designed the study. C.F. performed mouse experiments, intravital multiphoton imaging, flow cytometry, and image analysis. C.W. performed flow cytometry experiments. E.A.P. assisted in mouse husbandry. K.R.M. supervised the research and provided funding. C.F. and K.R.M. wrote the manuscript.

## Competing Interests

The authors declare no competing interests.

## Methods

### Mice

Mice were bred and maintained under specific pathogen-free conditions at Princeton University. All procedures were approved by the Institutional Animal Care and Use Committee (IACUC) and complied with relevant ethical regulations. Albino B6 (B6(Cg)-Tyrc-2J/J, Jax, 000058), *Rag2*⁻/⁻ (JAX 002216), *Csf1r*-EGFP (B6.Cg-Tg(Csf1r-EGFP)1Hume/J, JAX 018549), *Foxp3*-RFP (C57BL/6-Foxp3tm1Flv/J, JAX 008374), *Foxp3*-DTR (B6.129(Cg)-Foxp3tm3(HBEGF/GFP)Ayr/J, JAX 016958), and *Pdgfrα*-H2BGFP (C57BL/6-Pdgfratm1(H2B-GFP)Jrt/J, JAX 013148) mice were obtained from The Jackson Laboratory. *IL-10⁻/⁻* (JAX 002251) mice were obtained from the laboratory of J. Brooks II. All imaging and experimental manipulations were performed on mouse hind digits. Preparation of digits for intravital imaging was performed as described below. In brief, mice were anaesthetized with intraperitoneal injection of ketamine–xylazine (15 mg/ml and 1 mg/ml, respectively in PBS). After imaging, the mice were returned to their housing facility. For subsequent revisits, the same mice were processed again with injectable anesthesia. The hind digit regions were briefly cleaned with PBS pH 7.2, mounted onto a custom-made stage and a glass coverslip was placed directly against the digit tips. Anesthesia was maintained throughout the course of the experiment with vaporized isoflurane delivered by a nose cone. Mice from experimental and control groups were randomly selected from either sex for live imaging experiments. All experiments were repeated in at least three independent cohorts.

### Intravital multiphoton microscopy

P3 digit tips were visualized using a Leica multiphoton STELLARIS 8 DIVE system equipped with Discovery NX and Chameleon Vision II (Coherent) tunable Ti:Sapphire lasers. Imaging was performed through a water-immersion objective (N.A. 0.95, Leica) using a 600 Hz resonant scanner, with a 0.5 mm² field of view. Z-stacks were collected in 10 μm steps to a depth of approximately 300 μm, encompassing the epidermis, dermis, subcutaneous tissue, and P3 bone. Excitation wavelengths were 940 nm for GFP and 1,080 nm for RFP and second harmonic generation (SHG). Regions of interest (∼1.5 × 2 mm) were positioned on the P3 digit tip, guided by the SHG collagen signal from bone. All experiment were performed with at least three mice in each group.

### Image analysis

Raw image stacks were imported into Fiji (NIH) or Imaris (Oxford Instruments) for processing. Images are displayed as maximal projections of the full 300 μm optical stack. Large, tiled image stacks were acquired at consistent anatomical positions and aligned across time points in Imaris using all channels. Bone degradation area, regeneration area, and cell counts were quantified from total region of interest, blinded to experimenter to minimize observer bias.

### Digit amputation

Digit amputations were performed in 6–8-week-old mice as previously described^12^ . Analgesia and anesthesia were achieved by intraperitoneal injection of meloxicam (5 mg/kg) and a ketamine/xylazine mixture (15 mg/ml and 1 mg/ml, respectively, in PBS). Using sterile scalpels, the distal tip of the terminal phalanx (P3) of digits 2–4 of each hindlimb was removed to produce the regenerative (P3 R) condition, or the proximal P3 was amputated to produce the scarring (P3 NR) condition. Calcein (20 mg/kg; Sigma-Aldrich) and Alizarin red S (25 mg/kg; Sigma-Aldrich) were administered by intraperitoneal injection at DPA 0 to label mineralizing bone surfaces.

### Drug treatments and depletions

Treg depletion was achieved by intraperitoneal administration of diphtheria toxin (DT; Sigma-Aldrich, D0564; 30 ng/g body weight) in *Foxp3-*DTR mice every three days beginning at DPA −1. Control animals received equivalent volumes of PBS. CD4+ and CD8+ T cells were depleted by intraperitoneal injection of anti-mouse CD4 (clone GK1.5) and anti-mouse CD8α (clone 2.43) antibodies (200 μg each) or corresponding isotype controls at DPA −3, −1, 3, 7, 10, 14, and 17. For CXCR4 blockade, AMD3100 (Sigma-Aldrich; 15 μg/g) was administered by daily intraperitoneal injection beginning at DPA 0 through DPA 7.

### Mouse antibodies and flow cytometry

The following monoclonal antibodies were purchased from Thermo Fisher, BD Biosciences, BioLegend or Abcam: NK1.1 (PK136, 1:100), CD45 (30-F11, 1:300), CD4 (GK1.5, 1:100), CD4 (RM4-5, 1:100), CD11b (M1/70, 1:100), Ly6G (1A8, 1:250), Ly6C (HK1.4.rMAb, 1:250), CD8β (H35-17.2, 1:400), TCRβ (H57-597, 1:200) and CD3 (17A2, 1:100). Anti-mouse CD16/32 (BD Biosciences clone 2.4G2, 1:100) was used to block Fc receptors. DAPI (BD Biosciences, 564907, 1:5,000) was used to exclude dead cells.

P3 digit tips were harvested and finely minced into approximately 1–2 mm³ fragments. Tissue fragments were enzymatically digested in Liberase (280 μg/ml, Roche) at 37°C for 1 h with gentle pipetting every 15 min. The resulting cell suspension was filtered through a 70 μm strainer, pelleted at 400 × g for 5 min at 4°C, and resuspended in FACS buffer. Fc receptors were blocked with anti-mouse CD16/CD32 (2.4G2; BD Biosciences), and dead cells were excluded using DAPI (BD Biosciences). Stained cells were analyzed on a FACSymphony A3 (BD Biosciences). Data were acquired using FACSDiva software and analyzed with FlowJo v11.

### Micro-computed tomography

P3 digit tips were scanned using a Zeiss Xradia 630 Versa 3D X-ray microscope (Imaging and Analysis Center, Princeton University). Regions of interest corresponding to the distal P3 bone were manually segmented, and quantitative analyses of bone volume and bone length were performed blinded to experimenter.

### Statistical analysis

Data are expressed as mean ± SD or mean with paired lines. Student’s *t*-test was used for comparisons between two groups. One-way or two-way ANOVA with Tukey or Fisher post-hoc multiple comparisons tests was applied for three or more groups. *p* < 0.05 was considered significant. Statistical calculations were performed using Prism (GraphPad, USA).

### Reporting summary

Further information on research design is available in the Nature Portfolio Reporting Summary linked to this article.

## Data Availability

Source data are provided with this paper. All other data supporting the findings of this study are available from the corresponding author on reasonable request.

## Extended Data Figure Legends

**Extended Data Figure 1.**
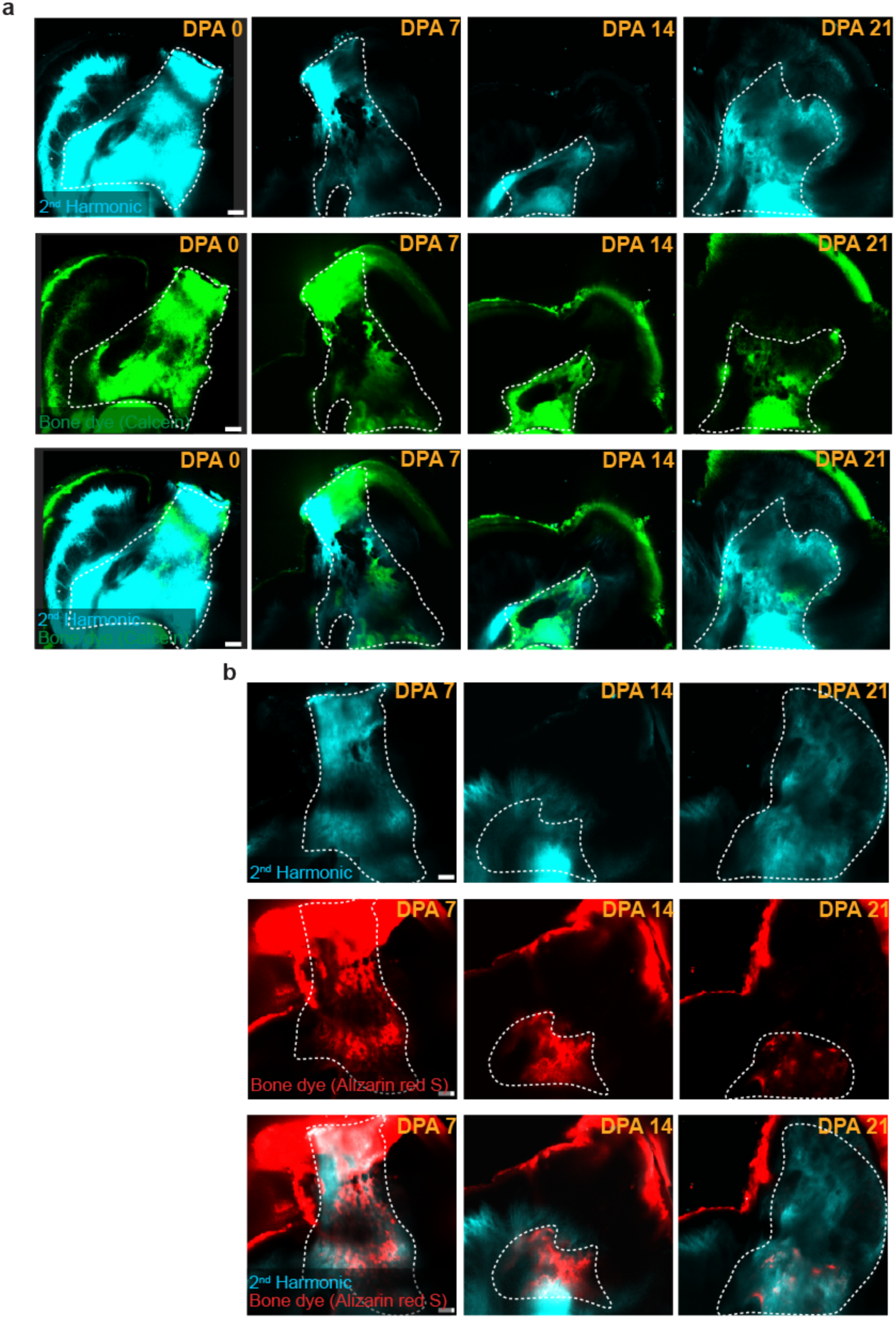
Validation of SHG as a longitudinal readout of bone remodeling. a, Representative serial two-photon images showing SHG (cyan, rows 1 and 3) and Calcein bone dye (green, row 2) at DPA 0, 7, 14 and 21 in P3 R digit tips, with merged images in row 3 demonstrating co-localization. Dashed white lines outline P3 bone. **b,** Representative serial two-photon images showing SHG (cyan, rows 1 and 3) and Alizarin red S bone dye (red, row 2) at DPA 7, 14, and 21 in P3 R digit tips. Dashed white lines outline P3 bone. Scale bars, 100 μm.

**Extended Data Figure 2.**
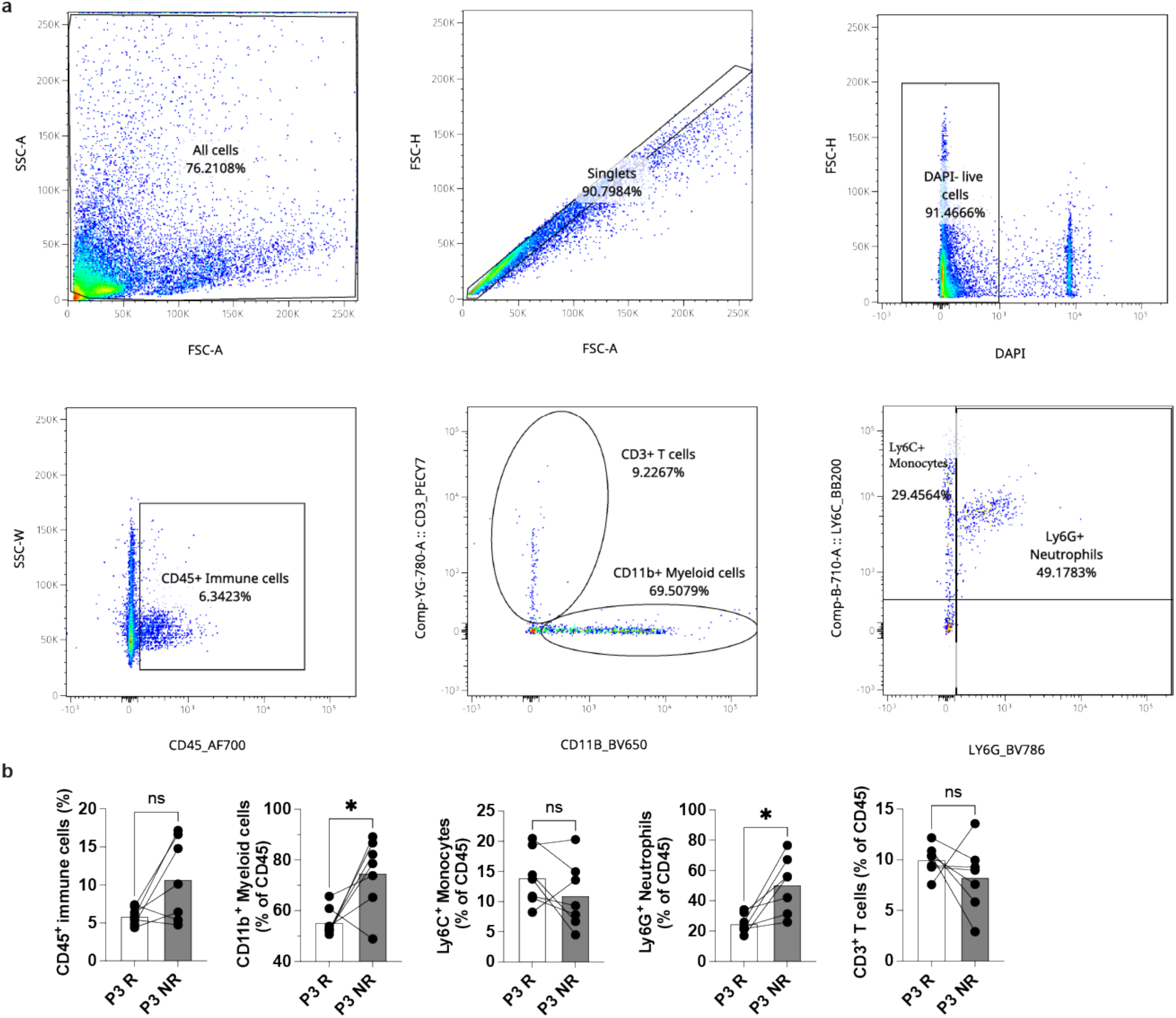
Immune cell profiling of P3 digit tips at DPA 7 by flow cytometry. a, Representative gating strategy for flow cytometry. **b**, Flow cytometric quantification of immune cell populations in P3 R and P3 NR digit tips at DPA 7, shown as paired data from the same mouse. CD45+ immune cells as a percentage of live cells; CD11b+ myeloid cells and CD3+ T cells as percentages of CD45+ cells; Ly6C+ monocytes and Ly6G+ neutrophils as percentages of CD45+ cells (n = 7 mice for each group, mean ± SD; *p < 0.05; ns; paired Student’s t-test).

**Extended Data Figure 3.**
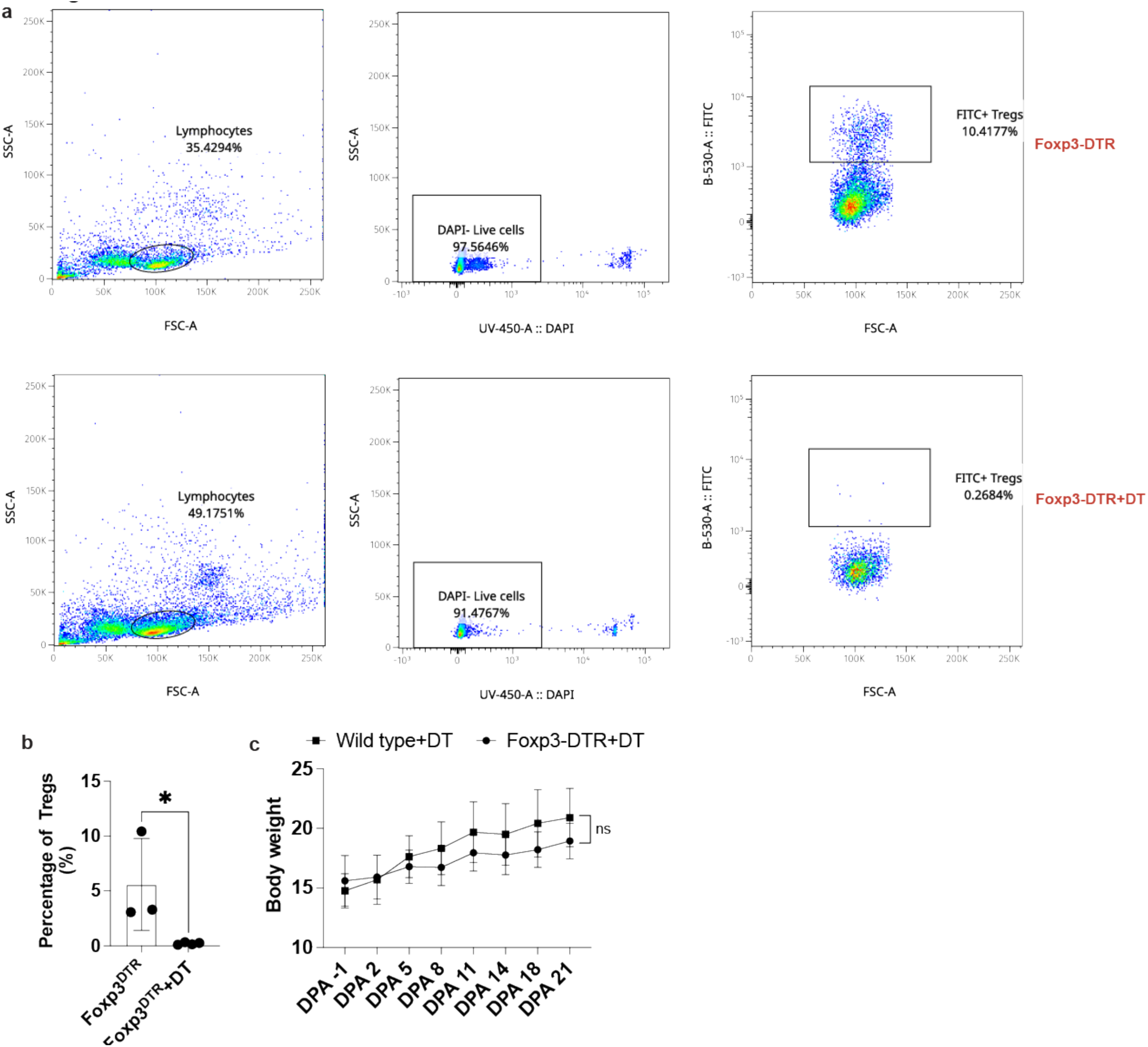
Validation of Treg depletion in *Foxp3-DTR* mice. a, Representative flow cytometry plots showing Foxp3+ cells (Tregs) in blood from *Foxp3-DTR* control mice (top) and from *Foxp3-DTR* + DT mice at DPA 28, demonstrating efficient depletion of circulating Tregs. **b,** Quantification of Treg percentage in blood at DPA 28 in *Foxp3-DTR* + DT (n = 3 mice) versus *Foxp3-DTR* control mice (n = 4 mice) (mean ± SD; *p < 0.05; unpaired Student’s t-test). **c,** Body weight of Wild-type + DT (n = 10 mice, block) and *Foxp3-DTR* + DT (n = 10 mice, spot) mice from DPA −1 through DPA 21 (mean ± SD; two-way ANOVA).

**Extended Data Figure 4.**
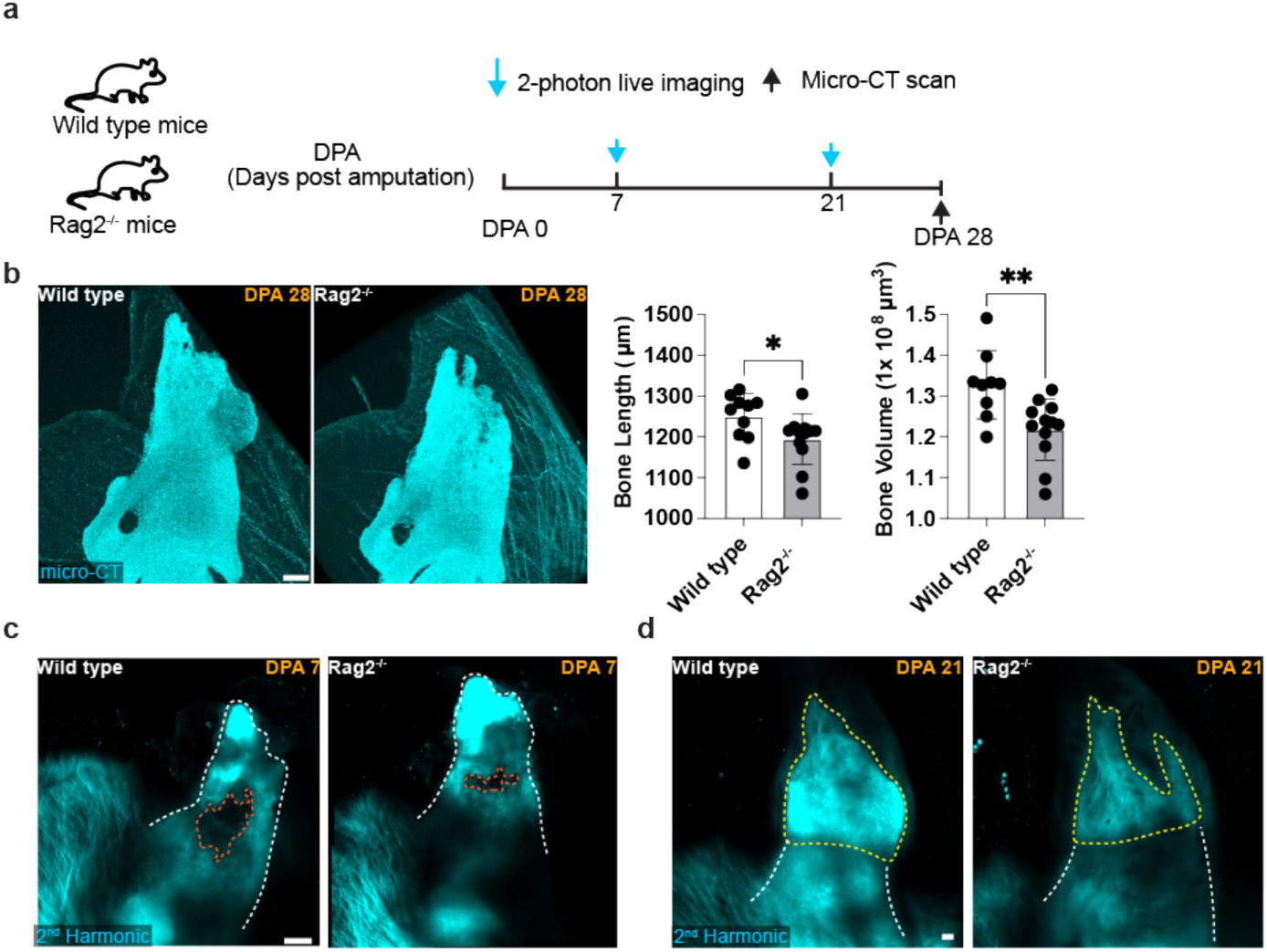
Lymphoid-deficient *Rag2*^-/-^ mice show impaired digit tip regeneration. a, Schematic timeline showing serial two-photon imaging at DPA 7 and 21 and micro-CT scan at DPA 28 in Wild-type and *Rag2*^-/-^ mice after distal P3 amputation. **b,** Representative micro-CT images at DPA 28 and quantification of P3 bone length (left) and bone volume (right) in Wild-type (n = 10 mice) versus *Rag2*^-/-^ mice (n = 10 mice) (mean ± SD; *p < 0.05, **p < 0.01; unpaired Student’s t-test). **c,** Representative two-photon SHG images at DPA 7 in Wild-type (left) and *Rag2*^-/-^ (right) mice. Dashed white lines outline P3 bone; dashed orange lines outline degradation area. **d,** Representative two-photon SHG images at DPA 21 in Wild-type (left) and *Rag2*^-/-^ (right) mice. Dashed white lines outline P3 bone; dashed yellow lines outline regeneration area. Scale bars, 150 μm.

**Extended Data Figure 5.**
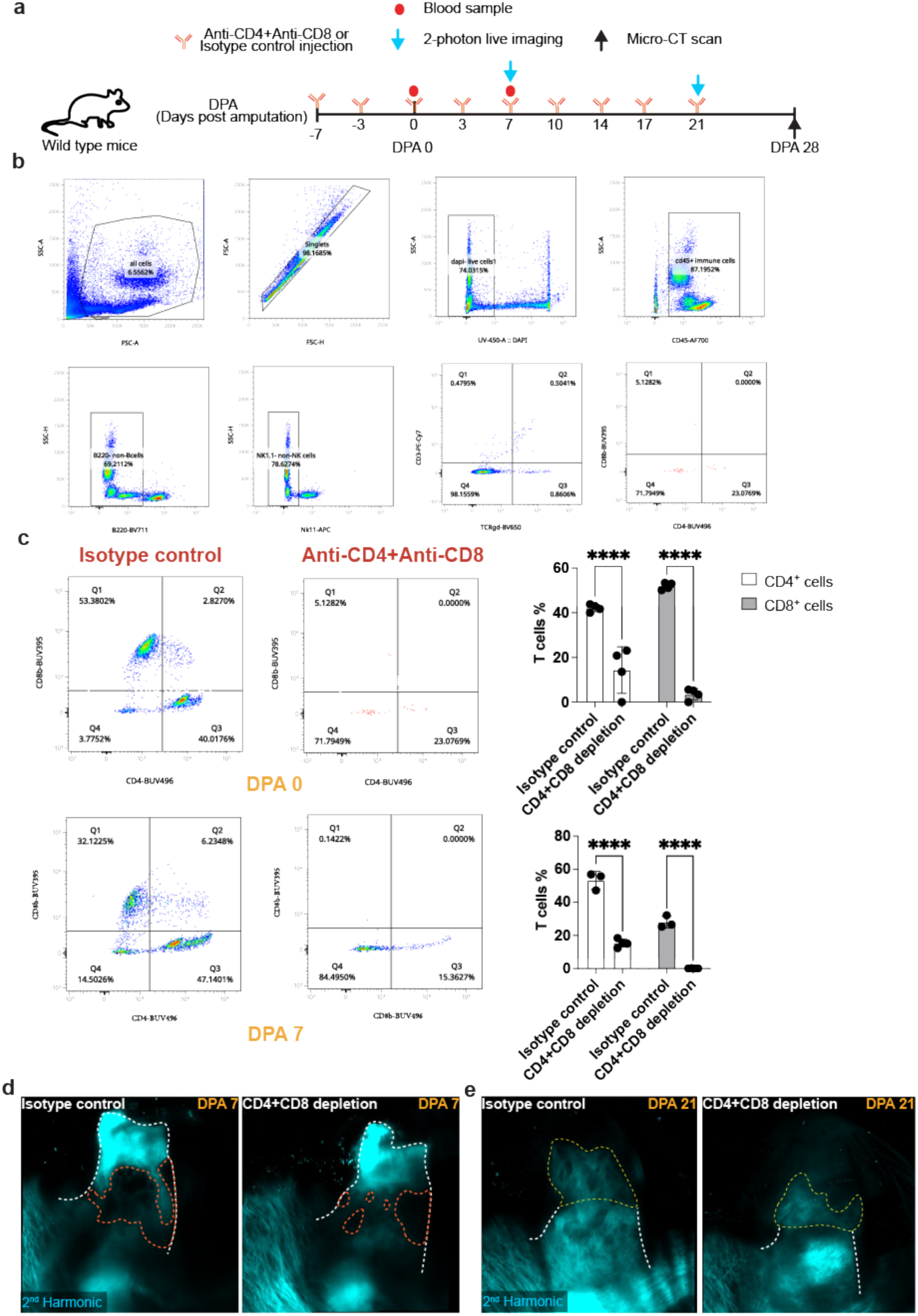
Anti-CD4 and anti-CD8 antibody depletion impairs digit tip regeneration. a, Schematic timeline of antibody injection schedule (anti-CD4 + anti-CD8 or isotype control, beginning at DPA −7), cheek bleeds for flow cytometric validation, two-photon imaging, and micro-CT. **b,** Representative gating strategy for flow cytometry. **c,** Representative flow cytometry plots and quantification of CD4+ and CD8+ T cells in peripheral blood at DPA 0 and DPA 7, confirming sustained depletion in anti-CD4 + anti-CD8-treated (n = 3-4 mice) versus isotype-control mice (n = 4 mice) (mean ± SD; ****p < 0.0001; unpaired Student’s t-test). **d,** Representative two-photon SHG images at DPA 7 in isotype-control (left) and anti-CD4 + anti-CD8-depleted (right) mice. Dashed white lines outline P3 bone; dashed orange lines outline degradation area. **e,** Representative two-photon SHG images at DPA 21 in isotype-control (left) and anti-CD4 + anti-CD8-depleted (right) mice. Dashed white lines outline P3 bone; dashed yellow lines outline regeneration area. bars, 150 μm.

**Extended Data Figure 6.**
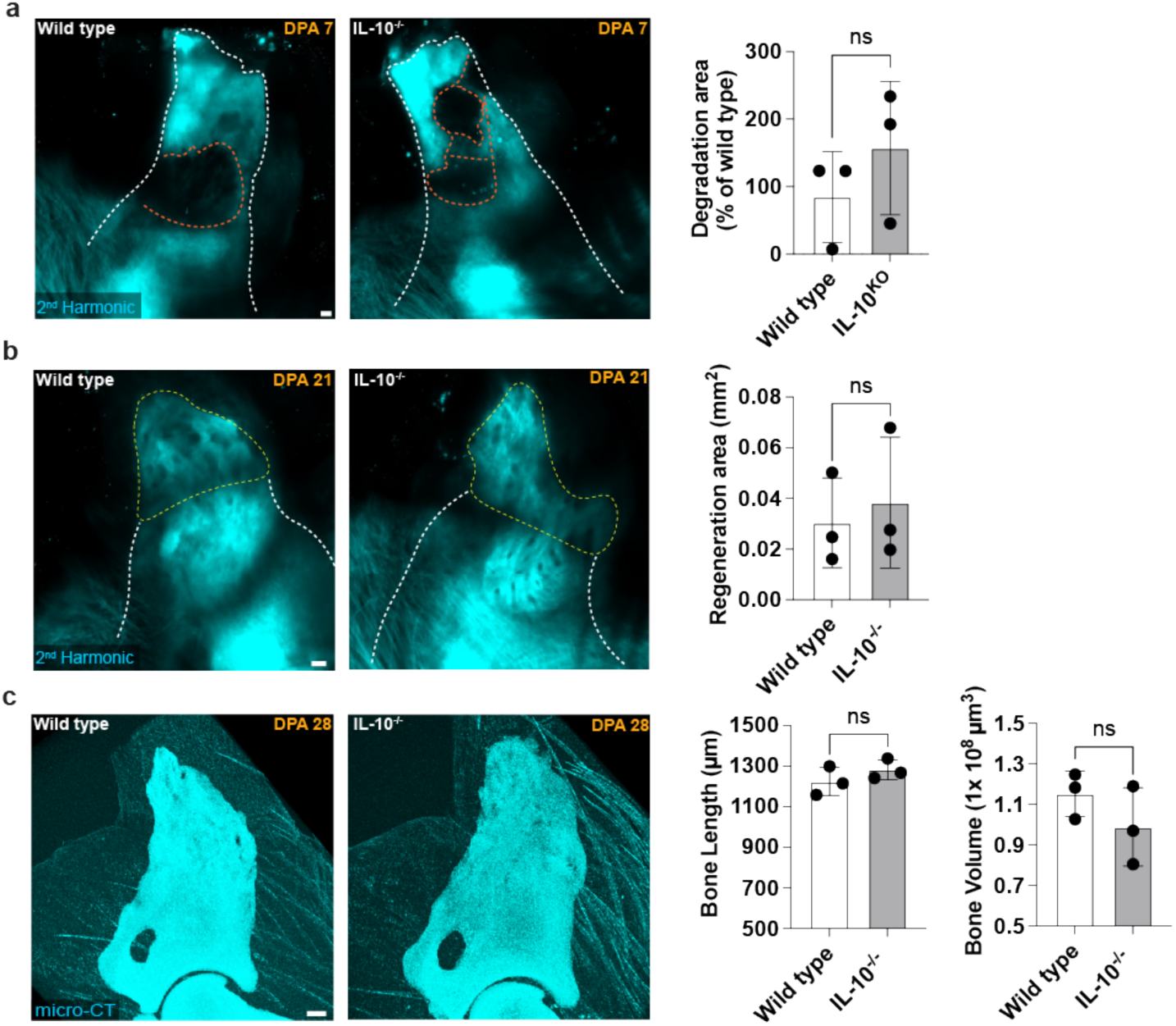
IL-10 deficiency does not impair digit tip regeneration. a, Representative two-photon SHG images at DPA 7 in Wild-type (left) and IL-10^-/-^ (right) mice, and quantification of bone degradation area in Wild-type (n = 3 mice) versus IL-10^-/-^ mice (n = 3 mice) (mean ± SD; ns; unpaired Student’s t-test). Dashed white lines outline P3 bone; dashed orange lines outline degradation area. **b,** Representative two-photon SHG images at DPA 21 and quantification of bone regeneration area in Wild-type (n = 3 mice) versus IL-10^-/-^ mice (n = 3 mice) (mean ± SD; ns; unpaired Student’s t-test). Dashed white lines outline P3 bone; dashed yellow line outline regeneration area. **c,** Representative micro-CT images at DPA 28 and quantification of P3 bone length (left) and bone volume (right) in Wild-type (n = 3 mice) versus IL-10^-/-^ mice (n = 3 mice) (mean ± SD; ns; unpaired Student’s t-test). Scale bars, 100 μm.

## Supplemental Video Legends

**Supplemental Video 1.** Two-photon images showing SHG bone signal (cyan) and Calcein (green) in P3 R digit tips at unamputated (left), DPA 7 (middle) and DPA 28 (right). Note Calcein and second-harmonic generation (SHG) signals reveal pre-existing bone (cyan + green) and newly formed bone (cyan). Note degradation site at DPA 7 and bone regrowth at DPA 21. Scale bar, 200µm.

**Supplemental Video 2.** 3D micro-CT reconstruction of the digit tip at DPA 7. Left: P3 R showing clear degradation site. Right: P3 NR with no visible degradation. Note: The image highlights the differential bone degradation between regenerating (P3R) and non-regenerating (P3NR) digit tips. Scale bar, 200µm.

**Supplemental Video 3.** Serial optical sections of digit tip in *Csf1r-eGFP; Foxp3-RFP* mice at DPA 7. Left: merged image showing osteoclasts / macrophages (green) and Tregs (red) localized in the degradation sites. Right: second-harmonic generation (SHG) signal. Note: This panel highlights the spatial association of osteoclasts and Tregs at sites of bone degradation during early regeneration. Scale bar, 100µm.

**Supplemental Video 4.** Serial optical sections of digit tip in *Pdgfrα-H2BGFP; Foxp3-RFP* mice at DPA 10. Left: merged image showing progenitor cells (green) and Tregs (red) precisely localized within the blastema pocket. Right: second-harmonic generation (SHG) signal. Note: The panel highlights the co-localization of progenitors and Tregs within the blastema microenvironment during active regeneration.

## Notes

### Competing Interest Statement

The authors have declared no competing interest.

